# Optimized ChIP-seq for Transcription Factors in Embryonic Mouse Brain

**DOI:** 10.64898/2026.11.13.766299

**Authors:** Alicia Li, Youngshin Lim, Ginam Cho

## Abstract

Chromatin immunoprecipitation followed by sequencing (ChIP-Seq) is widely used to identify genome-wide protein-chromatin interactions. However, obtaining reliable ChIP-seq data for transcription factors has been challenging for several reasons, including their low abundance, transient or weak DNA binding, and poor antibody performance. Here, we present an improved ChIP-Seq protocol for mapping transcription factor occupancy in embryonic mouse brains. We describe critical steps in tissue preparation, fixation and antigen unmasking steps, and chromatin fragmentation by prolonged sonication, specifically optimized for transcription factors. Our key innovation is the application of antigen retrieval to freshly fixed tissues, a technique commonly used in immunohistochemistry to unmask epitopes masked by fixation, in combination with slow fixation and extensive sonication to enhance ChIP-seq performance. This approach enhances epitope accessibility and expands the range of antibodies suitable for ChIP-seq, providing a robust strategy for mapping the genome-wide occupancy of transcription factors that are otherwise challenging to study using conventional ChIP-seq methods.

## INTRODUCTION

Transcription factors (TFs) are proteins that bind to specific DNA sequences (usually 6-12 bp) and regulate DNA transcription.^2^ They are particularly important during development, as they regulate how cells acquire specific identities and maintain them.^3^ Therefore, studying TF-DNA interaction is essential for understanding how TFs engage DNA regulatory elements to establish gene expression programs that drive cell fate specification and tissue development during embryogenesis.^4^ In addition, many human congenital diseases are attributed to genetic mutations of TFs, such as cortical malformation (ARX mutation), aniridia (PAX6 mutation), synpolydactyly (HOXD13 mutation), and schizencephaly (EMX2 mutation).^5^ Thus, there is a continual need to further elucidate the role of TFs in gene-regulatory mechanisms of disease.

ChIP-Seq is a powerful technique for identifying genome-wide protein-DNA interactions. Since its development, the technique has undergone continuous improvement, yet several challenges remain. One major limitation is the availability of suitable antibodies, as highly sensitive and specific antibodies are not readily for certain TFs.^2,6, 7^ Moreover, even if an antibody performs robustly in immunolabeling of fixed tissues, the same antibody may perform poorly in ChIP-seq. This limitation is compounded by the relatively low abundance of many TFs compared with histones and components of the core transcriptional machinery, which can reduce the amount of TF-DNA complexes available for ChIP enrichment.

Traditional ChIP-seq also requires relatively large amounts of starting material, which may not be available for certain cell or tissue types. ^8^ This presents an additional challenge when studying embryonic development, where the amount of tissue available at specific developmental stages can be limited. Thus, there remains a need for improved ChIP-seq protocols that increase the efficiency and sensitivity of TF detection.^9^ Recent modifications incorporating double cross-linking have been shown to improve ChIP-seq performance for TFs. However, these approaches still depend on the availability of high-quality ChIP-compatible antibodies and have been applied primarily to in vitro cell culture systems. ^9^

To address these limitations, we developed an adaptation of a commercially available ChIP-seq assay kit (Simplechip® Plus Sonication Chromatin IP Kit) by improving epitope accessibility for antibody binding in fixed tissue samples, potentially expanding the range of antibodies suitable for ChIP. The protocol incorporates optimized fixation, antigen retrieval, and chromatin fragmentation steps prior to immunoprecipitation.

The protocol described below was optimized for ChIP-seq analysis of embryonic mouse forebrain tissue collected at embryonic day 14.5 (E14.5). Library preparation and sequencing are not covered here; standard methods for preparing ChIP-seq libraries for next-generation sequencing can be used. For complete details on the use and execution of this protocol, please refer to Lim et al.^1^

## MATERIALS

| REAGENT or RESOURCE | SOURCE | IDENTIFIER |
|---|---|---|
| Antibodies |  |  |
| ARX antibody | R&D Technologies | AF7068; RRID: AB_10973178 |
| Critical commercial assays |  |  |
| Simplechip<sup>®</sup> Plus Sonication Chromatin IP Kit | Cell Signaling Technology | 56383 |
| QIAquick PCR Purification Kit | Qiagen | 28104 |
| Chemicals, peptides, and recombinant proteins |  |  |
| 16% Formaldehyde, Methanol-Free (PFA) | Cell Signaling Technology | 12606 |
| PBS (10X), pH 7.4 | Gibco | 70-011-044 |
| cOmplete EDTA-free Protease Inhibitor Cocktail (PIC) | Roche | 11697498001 |
| Antigen Unmasking Solution, Citric Acid Based | Vector Laboratories | H-3300-250 |
| Proteinase K (20 mg/ml) | Cell Signaling Technology | 10012 |
| RNAse A (10 mg/ml) | Cell Signaling Technology | 7013 |
| SYBR™ Safe DNA Gel Stain | Invitrogen | S33102 |
| UltraPure™ DNase/RNase-Free Distilled Water | Invitrogen | 10977015 |
| Hanks' Balanced Salt Solution (HBSS) |  |  |
| Deposited data |  |  |
| ARX ChIP-seq data | Young et. al.<sup>1</sup> | GSE274243 |
| Experimental models: Organisms/strains |  |  |
| CD1 mice | Charles River | 022 |
| Other |  |  |
| Water boiler | N/A |  |
| Dounce Tissue Grinder | N/A |  |
| Bioruptor UCD-200 Sonicator | Diagenode |  |
| Orbital shaker and rotator | N/A |  |
| Sample mixer | N/A |  |
| Tabletop centrifuge | N/A |  |
| 15 mL polypropylene tubes | N/A |  |
| P1000 wide-bore tips | N/A |  |
| Magnetic bead separation rack for 1.5 mL tubes | N/A |  |

### Mice

CD1 mice were used in this study. The day of the vaginal plug was designated as embryonic day E0.5.

### Antibodies

Typically, high-quality, ChIP-validated antibodies are required for ChIP-seq. We have modified this protocol to accommodate for ARX, an antibody that not been validated for ChIP-seq and that typically require antigen retrieval for tissue immunolabeling experiments. The amount of antibody used for immunoprecipitation may need to be optimized and experimentally determined. We recommend starting with 2 µg of antibody per immunoprecipitation experiment.

### Proteinase Inhibitor Cocktail (PIC, 200x)

Prepare a 200x PIC solution by thoroughly vortexing one cOmplete™ Protease Inhibitor Cocktail tablet in 250 µL Milli-Q water. Warm the solution in a 37°C water bath to help dissolve the tablet. The 200x PIC solution can be stored at -20°C for one week. This PIC 200x stock will be used in making the buffers to obtain a 1x PIC in the buffer.

### 1X ChIP Sonication Cell Lysis buffer

Prepare 2 mL of cell lysis buffer per sample according to SimpleChIP® Plus Sonication Chromatin IP Kit (per mL, add 0.5 mL 2X ChIP Sonication Cell Lysis Buffer, 0.5 mL water, 5 µL 200X PIC).

### ChIP Sonication Nuclear Lysis buffer

Prepare 1 mL of nuclear lysis buffer per sample by adding 5 µL of 200X PIC to 1 mL of SimpleChIP®’s

ChIP Sonication Nuclear Lysis buffer.

### Glycine Quenching reagent

Prepare 2 mL of glycine quenching reagent per sample by diluting the 10x Glycine solution provided in the SimpleChIP® Plus Sonication Chromatin IP Kit with UltraPure™ DNase/RNase-Free Distilled Water to a 1x Glycine solution. Make the quenching reagent right before use and use on the same day.

### PBS with PIC

Store PBS with PIC 1x at 4°C. Ideally, make the PBS with PIC and use it on the same day.

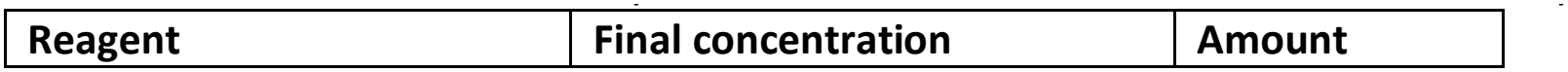
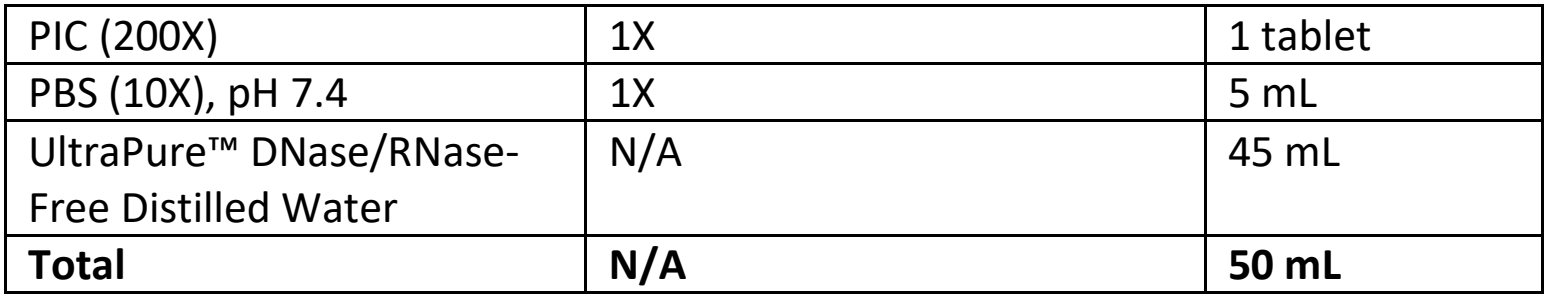

| **Reagent** | **Final concentration** | **Amount** |
|---|---|---|
| PIC (200X) | 1X | 1 tablet |
| PBS (10X), pH 7.4 | 1X | 5 mL |
| UltraPure™ DNase/RNase-Free Distilled Water | N/A | 45 mL |
| **Total** | **N/A** | **50 mL** |

### Paraformaldehyde fixing reagent (PFA 4%)

| **Reagent** | **Final concentration** | **Amount** |
|---|---|---|
| 16% Formaldehyde, Methanol-Free (PFA) | 4% | 10 mL |
| PBS (10X), pH 7.4 | 1x | 4 mL |
| UltraPure™ DNase/RNase-Free Distilled Water | N/A | 26 mL |
| **Total** | **N/A** | **40 mL** |

Keep PFA 4% on ice. Make PFA and use it on the same day.

**CRITICAL:** Formaldehyde is a known human carcinogen and toxic irritant. Wear personal protective equipment and prepare fixing reagent under a chemical hood.

### Antigen Retrieval (Ag) buffer

Thoroughly shake the Antigen Unmasking Solution, as it has a high salt concentration. Alternatively, warm the bottle in a 37°C water bath until no precipitates are seen inside the bottle. Prepare 3 mL of Ag buffer per sample and keep on ice. Use the solution on the same day.

| **Reagent** | **Final concentration** | **Amount** |
|---|---|---|
| Antigen Unmasking Solution (100X Citrate Buffer) | 1x | 0.03 mL |
| UltraPure™ DNase/RNase-Free Distilled Water | n/a | 2.97 mL |
| **Total** | **n/a** | **3 mL** |

## METHODS

### Embryonic Mouse Sample Preparation

This section describes the sample fixation process for one embryonic mouse brain. Other tissues of interest can also be harvested, but further optimization may be required. One well of a 12-well is used per brain sample.

#### Timing: [7h]

1. Prepare dissection workspace with the following:
  a. A bucket of ice.
  b. A 12-well plate with 2 mL of freshly prepared, ice-cold 4% PFA per well. Each sample will require one well. Place the well-plate on ice.
  c. Ice-cold HBSS.
  d. 2 6-cm petri dishes.
2. Fill the two petri dishes with ice-cold HBSS.

##### CRITICAL

It is important that the HBSS remains ice-cold to ensure tissue quality.

3. Humanely sacrifice the pregnant dam according to approved IACUC protocols and harvest the embryos. Place the embryonic sac in one petri dish.
4. Remove one embryo from the embryonic sac and place it in a second petri dish. Carefully dissect the brain of each embryonic mouse.
5. Place the dissected brain in one well of a 12-well.
6. After dissecting all the embryos, place the 12-well plate on an orbital shaker on its lowest setting and incubate at 4°C for 3 hours. **NOTE**: This fixation period is for a E14.5 mouse brain. Further optimization may be required for other tissue samples.
7. Prepare 3 mL of quenching reagent per tissue sample (see Materials and Equipment).
8. After incubation, remove the 4% PFA from the well using a transfer pipette, being careful to not touch the sample.
9. Add 3 mL of ice-cold PBS to the 12-well. Wash the brain sample by gently swirling the plate, then aspirate. Perform a total of 2 washes.
10. Quench the fixation reaction by adding 2 mL of quenching reagent to the 12-well. Place the plate at 4°C on an orbital shaker for 1 hour.
11. After incubation, remove the quenching reagent using a transfer pipette.
12. Repeat step 9 and perform a total of two washes using ice-cold PBS.
13. Aspirate the PBS and replace with fresh ice-cold PBS. Place the plate on a rocker for at least 45 minutes at 4°C before beginning the next section of the protocol.
14. In a petri dish, carefully dissect out the telencephalon. Confirm the mass of the forebrain sample. **NOTE:** Embryonic forebrains used in this protocol were about 150-200 mg. If another tissue sample is used, optimization of how much tissue sample is used for downstream may be required.
15. Place the dissected telencephalon into a 15-mL polypropylene tube. Keep the tube on ice until all samples have been dissected and ready to proceed to the next step.

### Epitope Unmasking By Antigen Retrieval

This step allows epitopes on the tissue that have previously been hidden due to crosslinking to be exposed. Since tissue is very soft after this step, take extra caution to not lose any sample during aspiration steps.

#### Timing: [30 min]

1. Prepare an ice bucket.
2. Prepare the required amount of Ag retrieval buffer and set it on ice.
3. Fill an electronic water boiler halfway with water and bring it to a boil.
4. While water is waiting to be boiled, add 3 mL of Ag buffer to the 15-mL polypropylene tube with the sample.

##### CRITICAL

Polypropylene tubes must be used because it can withstand high heat. Other plastics may melt while boiling and contaminate the sample.

5. Once the water comes to a boil, place the Falcon tube into the boiling water bath. Set a timer for twenty minutes after seeing the antigen retrieval buffer begin to boil.
6. After the 20-min antigen retrieval, remove the tube from the boiling-water bath and immediately place it on ice. Cool for 5 min, starting the timer when the tube is placed on ice. **NOTE:** Wear heat protective gloves, as the Falcon tube after boiling is very hot.
7. Carefully aspirate the Ag buffer, being careful to not aspirate the tissue sample.
8. Wash tissue samples by pipetting 5 mL of sterile PBS into the Falcon tube, gently swirl, then aspirate using a pipette.
9. Repeat Step 8 twice for a total of three washes.
10. Proceed immediately to the next step of the protocol.

### Chromatin Fragmentation Timing: [2.5h]

This section describes cell lysis, then nuclear lysis using sonication, resulting in fragmented chromatin of correct size to continue with immunoprecipitation.

1. Prepare 3 mL of 1X ChIP Sonication Cell Lysis Buffer with PIC and 1 mL ChIP Nuclear Lysis buffer (see reagent setup) per brain sample. Keep buffers on ice.
2. Ensure that the Dounce tissue grinder is clean by washing with 5 mL of DNase/RNase-Free distilled water, spraying it with 70% ethanol, then rinsing again with 5 mL of water.
3. Using a P1000 wide-bore pipette tip or a cut pipette tip, add 1 mL of 1X ChIP Sonication Cell Lysis buffer to the 15-mL polypropylene tube with the samples. Transfer one sample into a Dounce tissue grinder.
4. Using a tight-fitting pestle (Type A), disaggregate tissue pieces with 15 strokes. **NOTE:** By this stage, there should be no visible tissue chunks. Perform a couple more strokes if necessary.
5. Transfer cell suspension to a 1.5 mL Eppendorf tube and incubate the cell suspension on ice for ten minutes.
6. Rinse the Dounce tissue grinder (see step 2) prior to homogenizing a second sample.
7. Pellet the cells using a tabletop centrifuge and centrifuge the sample for ten seconds.
8. Remove supernatant and resuspend the pellet in 1 mL of cell lysis buffer. Incubate the sample on ice for 5 minutes.
9. Pellet the cells a second time using a tabletop centrifuge for ten seconds.
10. Remove supernatant and resuspend the pellet in 1 mL of nuclear lysis buffer.
11. Set up the Bioruptor UCD-200 (Diagenode; high power) to cycle at 30 seconds on, 30 sec off. Add ice and a small amount of water to the basin.
12. Fragment chromatin by sonication for a total elapsed time of 2 hours. Replenish the basin with ice and remove excess water every 15 minutes. **NOTE:** Sonication conditions are optimized using this specific sonicator; further optimization may need to be performed to obtain chromatin fragments consisting mostly around 150-300 bp.
13. Take 50 µL of unclarified cell lysate into a new 1.5 mL tube chromatin fragmentation analysis.
14. Clarify cell lysate by using a microcentrifuge and centrifuging at 21,000 x g for 10 minutes at 4°C.
15. Carefully transfer the supernatant into a new 1.5 mL tube. Take 50 µL of this supernatant for DNA concentration analysis.

Pause point: samples can be stored in -80°C until you are ready for the next step.

### Chromatin Fragmentation and Concentration Analysis Timing: [3h]

This step determines whether the sonication was successful and whether the DNA concentration is appropriate for immunoprecipitation.

1. To both the unclarified and clarified cell lysate, add 100 µL nuclease-free water, 6 µL 5M NaCl, and 2 µL RNAse A. Vortex to mix and incubate samples at 37°C for 30 minutes.
2. Add 2 µL of Proteinase K. Vortex to mix, then incubate samples at 65°C for 2 hours. **NOTE:** Proteinase K digestion can also be performed overnight at 37°C.
3. Purify DNA samples using Qiagen DNA purification spin columns (see DNA purification).
  a. Elute the unclarified cell lysate in 5 µL of EB and the clarified cell lysate in 30 µL of EB.
4. Using the unclarified cell lysate, determine DNA fragment size by electrophoresis on a 1% agarose gel. Most (>90%) of the fragmented chromatin should be around 150-300 bp.
5. Determine the DNA concentration of the clarified cell lysate. The DNA concentration was measured to be at least 50 ng/µL.

### Chromatin Immunoprecipitation (IP) Timing: [8h]

This section describes how to purify TF-DNA complexes and reverse the cross-linking.

1. For each IP, calculate the volume of fragmented chromatin containing 5 µg. Dilute with 1× ChIP buffer containing PIC, using at least four volumes of buffer per volume of chromatin (chromatin ≥1:4). Prepare at least 510 µL of diluted chromatin per IP: 500 µL for the IP and an additional 10 µL for the 2% input.
2. Reserve 10 µL of the diluted chromatin as the 2% input. This represents 100 ng, or 2% of the 5 µg allocated to the IP.
3. Transfer 500 µL of the remaining diluted chromatin to a 1.5 mL tube and add the immunoprecipitating antibody. We used 2 µg of antibody per IP; this amount may require optimization for each antibody.
4. Incubate the IP samples while rotating on a sample mixer at room temperature (25°C) for two hours.
5. Thoroughly resuspend the ChIP-Grade Protein G Magnetic Beads by gentle vortexing. Immediately add 30 µL of the beads to each IP reaction and incubate at room temperature (25°C) for two hours.
6. Right before the two hours is up, prepare the washes. Prepare 5 mL of low salt wash, 1 mL of high salt wash, and 150 µL of 1X ChIP Elution buffer per IP sample according to Simplechip® Plus Sonication Chromatin IP Kit.
  a. Bring 10X ChIP buffer to room temperature, then add 500 µL 10X ChIP Buffer to 4.5 mL water for low salt wash. For high salt wash, add 100 µL 10X ChIP Buffer and 70 µL 5M NaCl to 900 µL water.
7. Set a thermomixer to 65°C.
8. Briefly centrifuge the tubes and pellet the Protein G Magnetic Beads in each IP by placing the tubes in the magnetic separation rack. Wait 1 minute for the solution to clear, then remove the supernatant. **NOTE:** It is crucial to remove all residual supernatant to avoid contamination.
9. Add 1 mL of low salt wash to the IP. Invert to mix, then briefly vortex. Put sample on a hula shaker in the cold room at 4°C, then continue to rotate and incubate for 5 minutes.
10. Repeat low salt wash steps for a total of 5 times, being careful to remove all leftover supernatant in the tubes to avoid contamination.
11. After aspirating the last low salt wash, add 1 mL of the high salt wash to the IP. Put the tube on the hula shaker and incubate for 10 minutes at 4°C.
12. Pellet protein G Magnetic Beads by placing the tubes on the magnetic bead separator. Wait until the solution clears, then carefully remove the supernatant.
13. Add 150 µL of 1X ChIP Elution Buffer to the 2% input sample and to each IP sample.
14. Elute chromatin from the protein G magnetic beads for 30 minutes at 65°C while vortexing at 1,200 rpm.
15. Briefly centrifuge samples using a tabletop microcentrifuge for 5 seconds to collect evaporation from the tube cap.
16. Pellet the magnetic beads by placing the tubes on a magnetic separation rack and wait for the solution to clear.
17. Carefully transfer eluted chromatin in the supernatant to new 1.5 mL Eppendorf tubes.
18. To the 2% input sample and the eluted chromatin samples, reverse cross-links by adding 6 µL of 5M NaCl and 2 µL of Proteinase K (20 mg/mL). Gently vortex to mix and incubate for 2 hours at 65 °C.

**Pause point**: samples can be stored at -20°C for one week.

### DNA Purification Timing: [20 min]

This step removes any impurities in the DNA and prepares it for library amplification. The procedure below is adapted from Qiagen%s QIAquick PCR Purification Kit. All centrifuge steps are carried out at 17,900 x g (13,000 rpm).

1. Add ethanol to the PE buffer as indicated by the QIAquick PCR Purification kit.
2. Add 750 µL of PB Binding buffer to the IP sample (5 times the volume of IP sample) and 750 µL to the input sample and vortex briefly.
3. Add 450 µL of the sample to the QIAquick column.
4. Centrifuge for 30 seconds using a tabletop microcentrifuge at 17,900 x g (13,000 rpm).
5. Aspirate the liquid from the collection tube, then load the remaining sample onto the QIAquick column.
6. Centrifuge again for 30 seconds, then aspirate the liquid from the collection tube.
7. Add 750 µL of PE Buffer to the QIAquick column and centrifuge for 1 minute.
8. Aspirate the liquid from the collection tube, then centrifuge for ten seconds to remove residual wash buffer.
9. Remove the collection tube and place each QIAquick column onto a clean 1.5 mL microcentrifuge tube.
10. Add 30 µL of EB Buffer (10 mM Tris·Cl, pH 8.5) to the center of the QIAquick membrane and centrifuge the column for 1 minute.
11. Take the same 30 µL eluent and elute again a second time.
12. Determine the DNA concentration. The expected concentration is around 5-10 ng/µL.

**Pause point**: samples can be stored at -20°C until further analysis.

## RESULTS

### Optimization of embryonic brain fixation, antigen retrieval, and sonication conditions

Preserving the molecular state of chromatin and protein-chromatin interactions is essential for successful ChIP-seq analysis, particularly for TFs, which may exhibit transient or weak chromatin binding. Accordingly, fixation and antigen retrieval conditions must be carefully optimized to preserve TF-chromatin interactions while maintaining epitope accessibility. Proper chromatin fragmentation is also critical for efficient immunoprecipitation and high-resolution mapping of TF occupancy.

Inadequate fragmentation can reduce mapping resolution and increase background signal by allowing larger chromatin fragments containing additional protein-DNA interactions to be co-immunoprecipitated. We therefore optimized tissue crosslinking and chromatin sonication conditions by testing different fixation and sonication durations (Figure 1), while maintaining consistent antigen-unmasking conditions across all samples.

**Figure 1.**
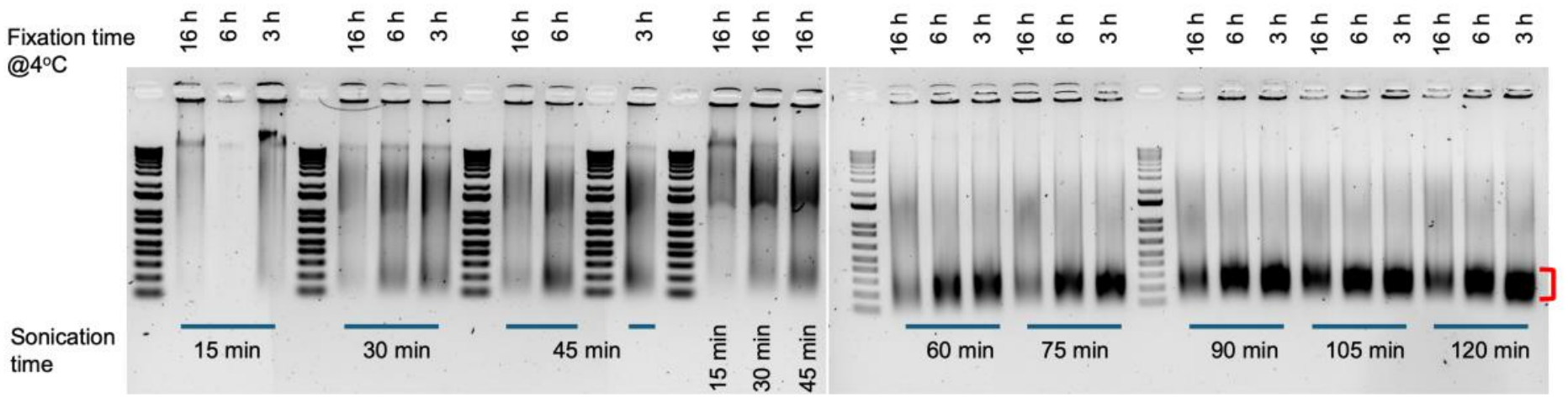
Representative agarose gel image of DNAs in isolated nuclei from mouse embryonic brains (E14.5) fixed with 4% paraformaldehyde at 4°C for overnight (16hr), 6 hr, or 3 hr, and fragmented by varying sonication time ranging from 15 min to 12 min. Top numbers indicate fixation time and bottom numbers indicate sonication duration time (30 s on; 30 s off). Red bracket indicates ideal fragment size (150-300 bp) of DNAs.

E14.5 mouse brains were fixed in 4% paraformaldehyde for 3h, 6h, or 16 h duration at 4°C. Following antigen unmasking by boiling tissues in a citrate-containing antigen retrieval solution for 20 minutes, a series of sonication conditions were tested using a Bioruptor UCD-200 at high power with 30 seconds on/30 seconds off cycles for 15, 30, 45, 60, 75, 90, 105, or 120 minutes. Among the conditions tested, 3 hour of tissue cross-linking with 4% paraformaldehyde followed by 120 minutes of sonication using 30-s on/30-s off cycles yielded optimal chromatin fragmentation, with the majority of fragments ranging from approximately 150-300 bp (Figure 1).

### ARX ChIP-seq as an example

We have tested this protocol using Aristaless-related homeobox (ARX), a key transcription factor for cortical interneuron development and function. ChIP-seq results from Young et. al^1^ shows nearly 50% of annotated transcripts being within 2kb of transcript start sites (TSS) (Figure 2). Majority of the annotated peaks were within promoter and exonic regions (Figure 3). An example of ARX ChIP-seq signal at the *Lmo1* locus, a known binding target of ARX, is provided (Figure 4).

**Figure 2.**
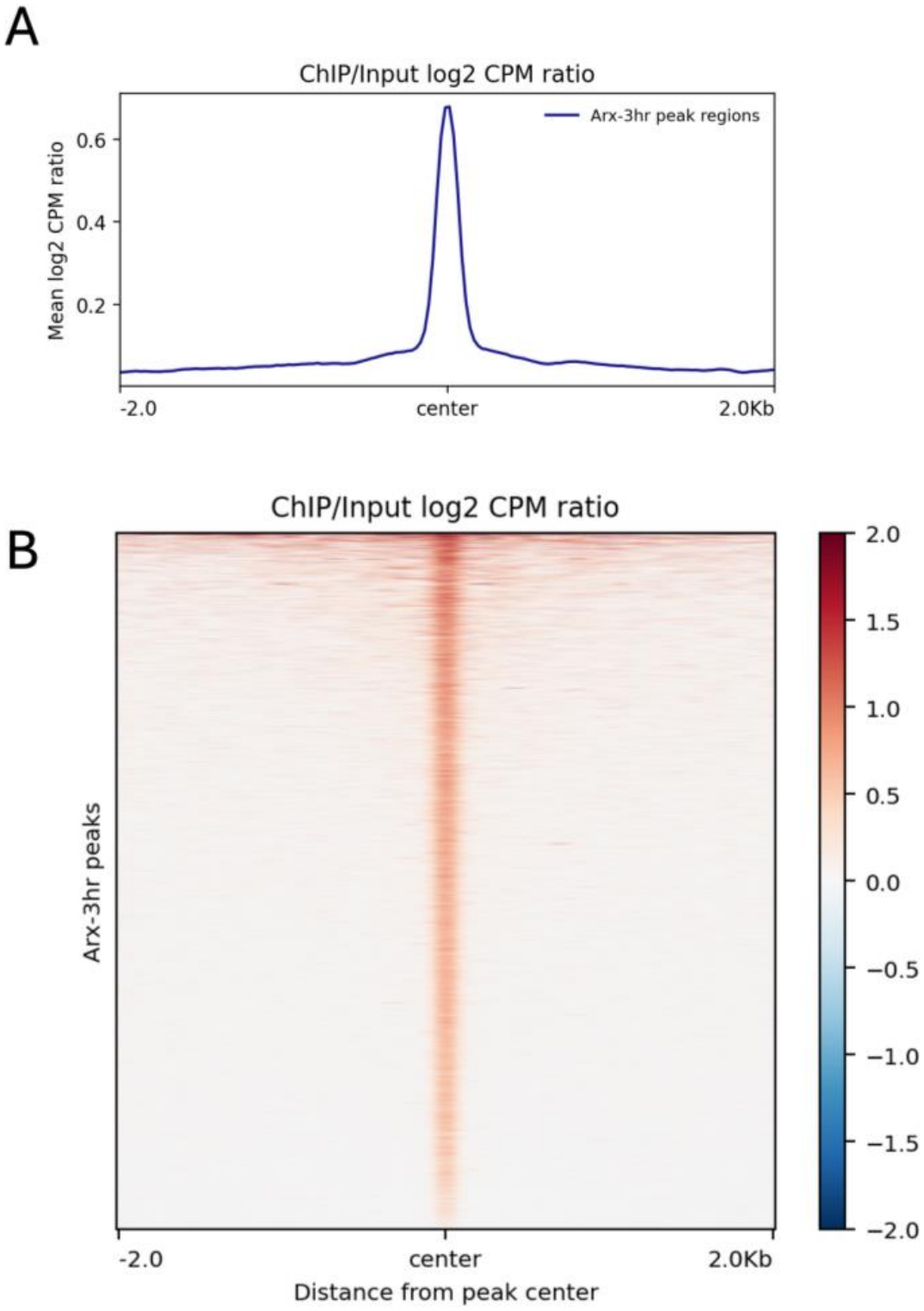
ARX ChIP-seq signal enriched at ARX peak regions. (A) Average profile of the ChIP/Input signal across ARX peak regions (n = [12,964] peaks). The signal is the log2 ratio of counts-per-million (CPM)-normalized ARX ChIP reads to input reads, shown ±2 kb from the peak center in [10]-bp bins. (B) Heatmap of the same signal, with one row per peak and peaks sorted by decreasing signal intensity. The color scale represents the log2 ChIP/Input ratio (red, enriched relative to input; blue, depleted relative to input). Data are from E14.5 mouse embryonic brains collected after 3hr of fixation at 4°C with 4% paraformaldehyde and normalized to the matched input. Peaks were called with [MACS2, q < 0.05], and signal tracks were computed with deepTools bamCompare.

**Figure 3.**
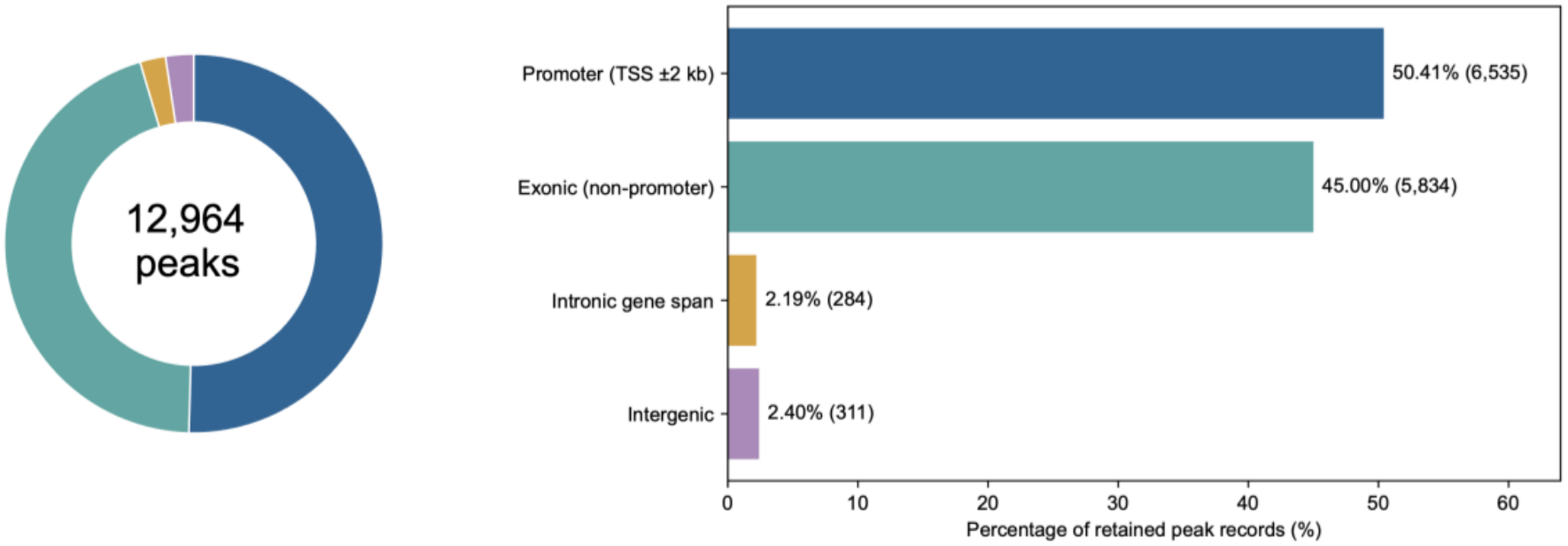
Genomic distribution of ARX ChIP-seq peaks. Peaks were annotated using GENCODE M25 gene annotations for the mm10 mouse genome assembly and classified according to their genomic location. To ensure mutually exclusive assignment, peaks overlapping multiple genomic features were assigned using the following priority: exclusive overlap priority is promoter > exon > gene span > intergenic. Promoter regions were defined as regions within any within 2kb of the annotation transcript TSS (Transcription Start Site). The distribution shows the proportion of ARX peaks assigned to each genomic category. Data were collected from E14.5 mouse brains fixed with 4% paraformaldehyde at 4°C for 3 hrs and sonicated for 120 min (30 s on; 30 s off).

**Figure 4.**
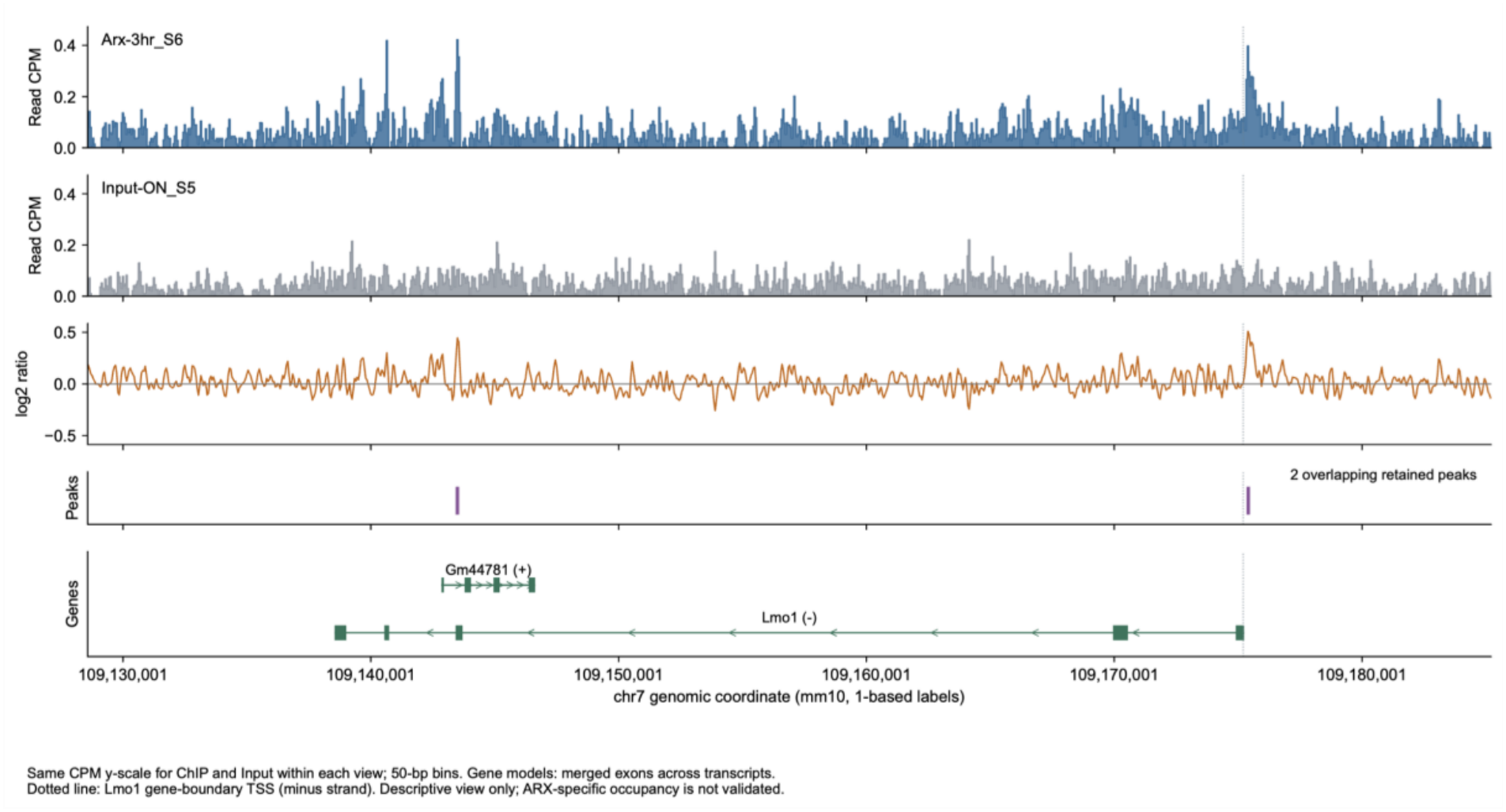
An example of ARX ChIP-seq signal (at the *Lmo1* locus). Genome browser view of the *Lmo1* gene body ±10 kb (chr7:109,128,572–109,185,207, mm10). Tracks from top to bottom: **(i)** ARX ChIP signal and **(ii)** matched input, both shown as read counts per million (CPM) in 50-bp bins on the same y-axis scale; **(iii)** log2 ratio of ChIP to input CPM; **(iv)** retained ARX peaks (n = 2), one in an *Lmo1* intron and one at the *Lmo1* transcription start site; **(v)** gene models (exons merged across transcripts; arrows show the strand), with *Lmo1* on the minus strand and *Gm44781* on the plus strand. The dotted vertical line marks the *Lmo1* TSS. Data were collected from E14.5 mouse brains fixed with 4% paraformaldehyde at 4°C for 3 hrs and sonicated for 120 min (30 s on; 30 s off)

## DISCUSSION

Conventional ChIP-seq protocols typically use formaldehyde fixation for 1 hour at room temperature. However, we utilized a slower fixation process—paraformaldehyde fixation for 3 hours at 4°C, to preserve the molecular state of the chromatin and the delicate nature of TF-chromatin binding. In antigen retrieval step, the heating (boiling) process partially reverses paraformaldehyde-induced cross-links formed during fixation, while citrate-based antigen retrieval promotes the restoration of epitope accessibility, thereby enhancing antibody recognition. Prolonged sonication step with total duration of 120 minutes for chromatin fragmentation in this protocol yielded optimal chromatin fragmentation.

This protocol has been successfully used for ARX ChIP-seq in embryonic mouse forebrains and may also be applicable to other transcription factors for which ChIP-grade quality antibodies are not available. However, paraformaldehyde fixation and sonication conditions may need to be optimized depending on the type and sonicator used.

Some limitations of this protocol are that the antigen retrieval step may not fully resolve ChIP-seq%s limitation of capturing indirect DNA-protein interactions. While it does enhance epitope availability, there may still be some binding targets that are unable to be pulled down via immunoprecipitation. Moreover, excessive cross-link reversal by antigen retrieval may disrupt protein-DNA cross-links (as well as protein-protein cross-links), therefore reducing the recovery of TF-associated DNA fragments, including directly bound target regions. This could result in weaker ChIP-seq enrichment and reduce peak signal.

In conclusion, we developed an optimized ChIP-seq methodology that addresses key challenges associated with profiling transcription factors in embryonic mouse brain tissue, including low TF abundance, fixation-induced epitope masking, limited antibody performance, and restricted tissue availability. By integrating optimized tissue fixation, antigen retrieval, and chromatin fragmentation, this approach improves epitope accessibility while preserving TF–chromatin interactions and generating chromatin fragments suitable for high-resolution analysis. Using ARX as a model neurodevelopmental TF, we demonstrate that this methodology enables robust mapping of TF occupancy in embryonic brain tissue. Importantly, the incorporation of antigen retrieval may expand the range of antibodies suitable for ChIP-seq, including antibodies that perform well in fixed tissues but have limited utility in conventional ChIP protocols. This methodology therefore provides a practical framework for investigating TF–chromatin interactions during neural development and may facilitate genome-wide studies of transcription factors that have previously been difficult to interrogate by ChIP-seq.

## LEAD CONTACT

Further information and requests for resources and reagents should be directed to and will be fulfilled by the Lead Contact, Ginam Cho.

## ACKNOWLEDGEMENTS

This work was supported by NIH NINDS grants R01 NS100007 and R01 NS113516. During the preparation of this manuscript, ChatGPT (GPT-6, ChatGPT for Healthcare Workspace) was used for the purposes of text editing, such as grammar, spelling, punctuation, or language improvement. The authors have reviewed and edited the output and take full responsibility for the content of this publication.

## AUTHOR CONTRIBUTIONS

G.C. and Y.L. conceptualized and supervised the study. G.C. performed the initial experiments, analyzed the data, and prepared the figures, and A.L. performed additional experiments. A.L. and Y.L. wrote the manuscript together and G.C. participated in manuscript revision.

